# Rhizosphere-colonizing bacteria persist in the protist microbiome

**DOI:** 10.1101/2025.01.16.633413

**Authors:** Stephen. J Taerum, Ravikumar R. Patel, Justin E. Alamo, Daniel Gage, Blaire Steven, Lindsay R. Triplett

## Abstract

Soils contain diverse predatory protists that affect the abundance and behavior of rhizosphere bacteria, including bacteria that may benefit plant health. Protists also harbor their own bacterial microbiomes, including transient and extracellular associates, but it is not known whether the protist microbiome affects the plant rhizosphere. To address this question, we profiled the bacterial microbiomes of eight evolutionarily diverse rhizosphere protist isolates after two years of continuous laboratory culture. We then compared the protist culture microbiomes to maize rhizosphere communities six weeks after protist inoculation. Introduction of protists enriched 13 protist-associated bacterial amplicon sequence variants (ASVs) in the rhizosphere, which comprised ∼10% of the rhizosphere bacterial community. Additional bacterial ASVs ranked highly in abundance in both rhizosphere and protist microbiomes; together, a median 47% of the protist microbiome was enriched or abundant in the rhizosphere. Inoculation with some of the protist cultures positively affected root biomass traits, but a protist mixture had no effect, indicating that the impact of protist holobionts on plant growth is context-dependent. Isolates of protist-associated bacteria had both positive and negative effects on protist growth in culture, suggesting that the bacteria use multiple strategies to survive in proximity to predators. This study demonstrates that even after prolonged laboratory culture, evolutionarily diverse rhizosphere protists host bacterial microbiomes dominated by plant-colonizing bacteria that impact the rhizosphere microbiome after inoculation. The findings suggest that protists may contribute to the rhizosphere as part of the soil microbial seedbank, and identify bacterial groups that may be important to the plant-protist interaction.

**Importance:** Understanding the impact of predatory protists on the plant microbiome will be essential to deploy protists in sustainable agriculture. This study shows that eight rhizosphere protist isolates hosted diverse and distinct bacterial communities, and that a large proportion of these bacteria could be found colonizing the maize root environment long after protists were introduced. This study demonstrates that maize rhizosphere bacteria can persist for years in the protist microbiome, indicating that extremely diverse eukaryotes could help select and maintain rhizosphere bacteria in the absence of the plant.

## Introduction

Microbial eukaryotes including fungi, nematodes, and protists are integral to the plant microbiome. Single-celled protists-eukaryotes that cannot be classified as plant, animal, or fungus-are ubiquitous in soil, and are often predators of bacteria and fungi (1). Protists may stimulate plant growth, and this was historically often attributed to the stimulation of nutrient cycling (2). Consensus has grown that predators also benefit plant growth and disease resilience through selective predation. Culling prey bacteria can increase the competitive fitness of predation-resistant bacteria, including plant growth promoting strains of *Bacillus*, *Sphingomonas, Pseudomonas*, and *Azospirillum* (3). Although few species of rhizosphere bacteria are known to benefit from protists, network association studies have identified hundreds of positive correlations among protist and bacterial groups in the rhizosphere (4–6). This suggests that a much wider diversity of rhizosphere bacteria may have positive ecological associations with predators than is currently established. Given that correlations could arise from niche constraints, environmental factors, or random chance (7), systematic studies are needed to understand which rhizosphere-colonizing bacteria are positively selected by protists.

Rhizosphere eukaryotes interact with bacteria, but they also host their own bacterial microbiomes that shape their interaction with plants. For example, nematode endosymbionts detoxify plant secondary metabolites, endofungal *Rhizobium* and *Burkholderia* spp. promote plant beneficial and pathogenic fungi, and *Fusarium*-colonizing *Pantoea* inhibit virulence by producing antifungal compounds (8–11). Highly cosmopolitan multi-kingdom colonists, including some strains of *Bacillus* and *Pseudomonas* bacteria, may even be transferred from fungi or nematodes to plants (12, 13). Protists host intracellular and extracellular bacterial communities through a wide range of mutualistic and antagonistic mechanisms (14). The model amoebae *Dictyostelium discoideum* hosts both prey and predation-resistant bacteria, which may be obligate or transient residents (15). Green algae microbiomes parallel those of plant roots, and include many bacteria that can colonize and benefit both organisms (16). No studies to our knowledge have characterized bacteria associated with other groups of rhizosphere protists, but an understanding of such communities could point to synergisms that benefit the plant. For example, nitrogen-fixing *Sinorhizobium meliloti* is a transient resident of ciliate cells, and association with protists increases the bacteria’s distribution and nodulation capacity (17, 18).

We previously observed that inoculating plants with a defined protist consortium resulted in a rhizosphere bacterial community composed of bacteria similar to those in the protist inoculum (19). Protist inoculation also alleviated a negative growth effect of field soil bacteria, and this impact was dependent on bacteria in the protist culture. These findings led us to hypothesize that the protist microbiome could help establish a beneficial rhizosphere microbiome. In this study, we identify bacterial taxa found in long-term culture with rhizosphere protist isolates. (Although the nature of their interaction with protists is not known, we term these “protist-associated bacteria” for brevity). Second, we tracked abundance of those bacterial taxa after protist inoculation onto plants. We asked, 1) Are protist-associated bacteria enriched in the rhizosphere after protist inoculation? 2) Do the microbiome effects of protist inoculation vary among different protists? 3) Is bacterial establishment in the rhizosphere affected by protist inoculum diversity? Finally, 4) how do protist-associated bacteria affect protist growth? We find that bacterial taxa associated with evolutionarily diverse protists can colonize the rhizosphere, suggesting that protists could affect rhizosphere health through shared microbiota.

## Results

### Characterization of bacteria associated with rhizosphere protist isolates

Protists were previously isolated from independent maize root samples at two sites (20), and maintained in laboratory culture for over two years without addition of live bacteria. Eight protist isolates were selected for microbiome characterization, of which five are known to colonize the maize rhizosphere after inoculation to roots (*Colpoda*, *Cercomonas, Ochromonas*, *Thaumatomonas*, and Chrysophyceae sp., Table S1) (19). Three isolates were selected to represent the groups Amoebozoa and Rhizaria (*Flamella* and *Nuclearia* sp.), and *Allapsa*, a genus with significance to plant health (5). Sequence profiling of 16S rRNA genes in twice-washed protist cultures identified 30 to 75 bacterial genera in each protist microbiome (Table S1). Protist culture microbiomes had lower Shannon diversity indices than bacteria cultured from bulk and rhizosphere soils collected at the original protist isolation sites, which we sequenced concurrently and used as bacterial inoculum for plants (Table S1).

We identified the 30 most abundant bacterial genera, belonging to 15 orders, across inoculum samples (henceforth referred to as “dominant genera”, Fig. 1). While each microbiome was distinct, there were five core dominant genera detected in all protist microbiomes (*Pseudomonas*, *Variovorax*, *Sphingomonas*, *Reyranella*, and *Bradyrhizobium*), and eight additional genera represented in six of eight communities. However, there were no core ASVs of *Sphingomonas* or *Reyranella*, and two core ASVs of *Variovorax* (Table S2), showing that the core genera include bacterial strain diversity. Members of Burkholderiales and Hyphomicrobiales were ubiquitously abundant in the protist microbiomes; nine of the dominant bacterial genera represented these two orders. The results indicate that while each protist culture enriched a distinct subset of bacteria after isolation from rhizosphere soil, certain bacterial taxa broadly persisted in and dominated microbiomes of diverse rhizosphere protists.

**Figure 1:**
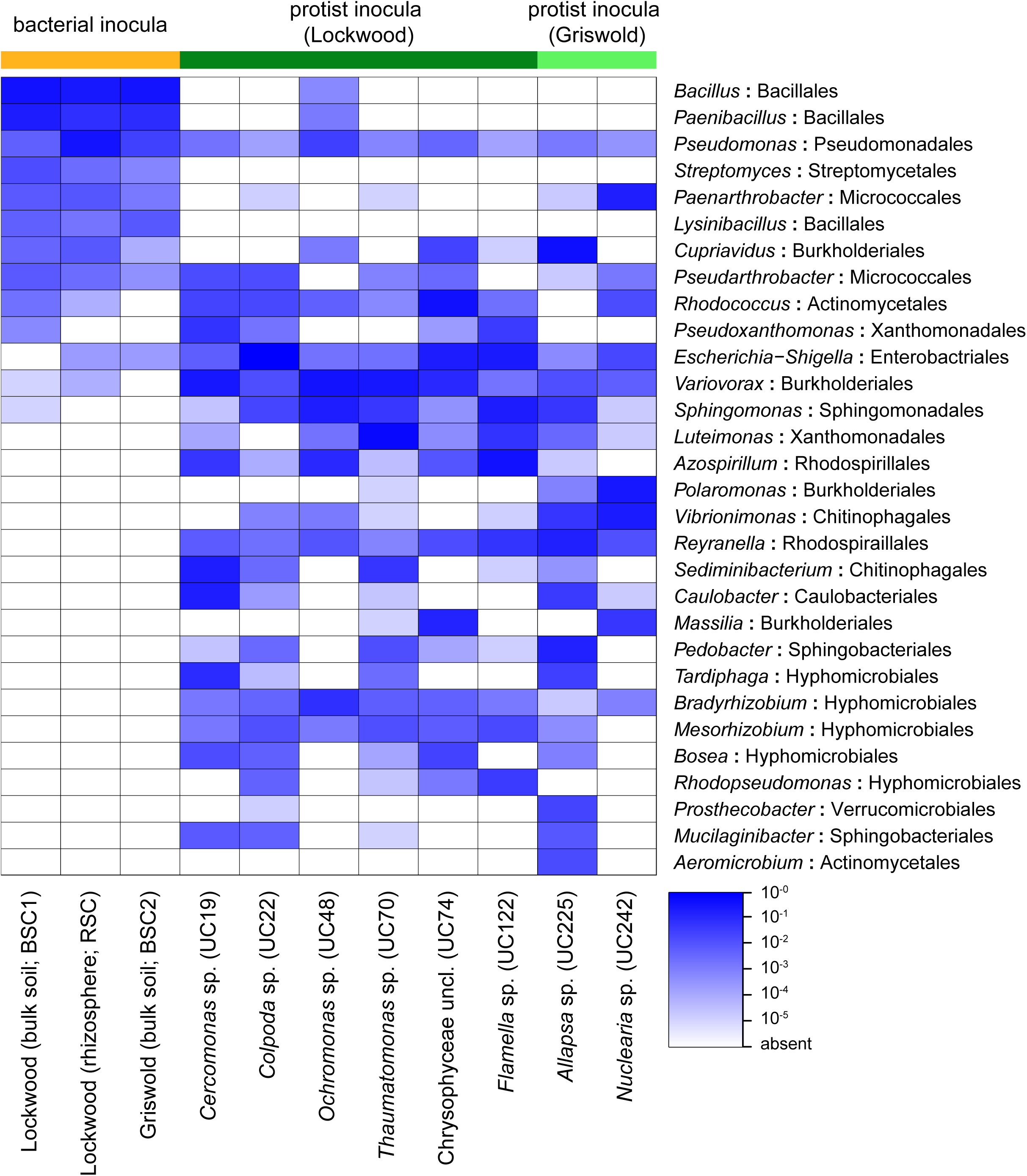
Bacterial microbiome composition of the protist-free bacterial inocula and the eight protist isolates from the maize rhizosphere. Genera having the highest mean abundance across samples are shown. Bacterial order is indicated after the colon. Shade corresponds to proportional abundance in the culture sample.

### Establishment of protists and protist-associated bacteria in the rhizosphere

To study the fate of protist-associated bacteria in the maize rhizosphere, we inoculated maize with the eight protist cultures singly and in mixture, and co-inoculated these with a bulk soil bacterial community cultured from our Lockwood Farm field site (BSC1, Fig. 2). To determine the effects of the bacterial community, additional plants were inoculated with bulk soil communities from Griswold (BSC2) or a rhizosphere soil community from the Lockwood site (RSC, Fig. 2), and co-inoculated with the eight--protist mixture. Rhizosphere microbial communities were profiled by amplicon sequencing of the 16S and 18S rRNA genes after six weeks.

**Figure 2:**
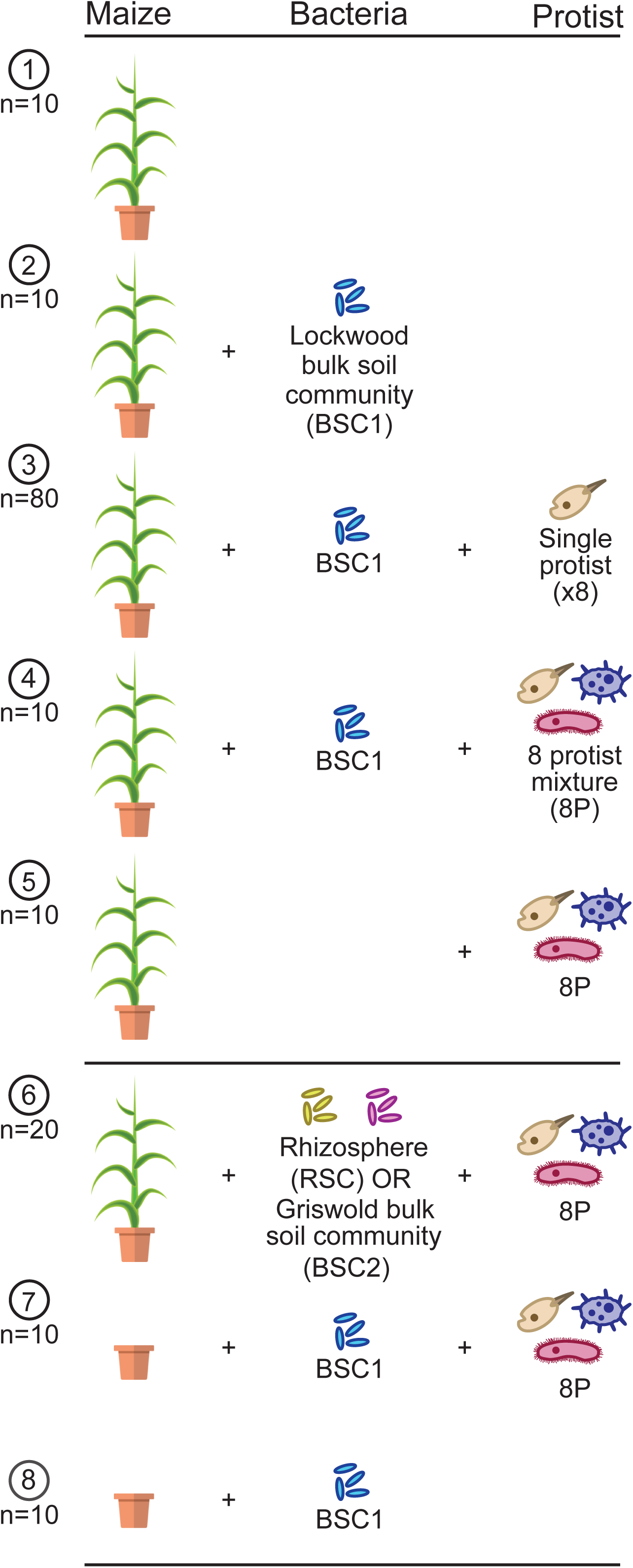
Experimental design showing eight combinations of protist and bacterial treatments. Numbers of planted replicates are listed under each circled number. Six plants failed to grow and eleven 16S rhizosphere libraries had fewer than 2000 reads, so these samples were removed from the study.

### *Colpoda* sp., but not other inoculated protists, were consistently abundant after six weeks

ASVs corresponding to four of the eight inoculated protists were detected in the rhizosphere samples after six weeks (Table S3); all were found to establish in the maize rhizosphere three weeks after inoculation in our previous growth chamber study: *Cercomonas* sp. (UC19), *Ochromonas* sp. (UC48), *Thaumatomonas* sp. (UC70), and *Colpoda* sp. (UC22) (19). In this study, the former three protists were detected inconsistently at six weeks, and at low read frequency (<0.01% of eukaryotic reads). In contrast, the *Colpoda* isolate ASVs consistently dominated protist sequence in inoculated pots, comprising an average of 20% of the eukaryotic reads (Table S3). Maize protist communities are highly temporally dynamic (21), this result suggests that *Colpoda sp*. is uniquely persistent as a plant inoculant.

Bacterial communities were highly variable within each bacterial treatment, and bacterial inoculation did not significantly affect rhizosphere diversity or plant biomass (Fig. S1). The lack of effect of bacterial inoculation suggested that bacteria from non-inoculum sources, such as the open greenhouse environment or maize seed, primarily determined the rhizosphere bacterial community composition.

### The maize rhizosphere consistently recruited specific protist-associated bacteria

Thirteen bacterial ASVs were significantly enriched in the rhizosphere after inoculation with individual protist cultures, even when the corresponding protists were not detected (Fig. 3A, Fig. S2). Bacterial taxa enriched in the rhizosphere typically comprised greater than 0.5% of the bacterial community in the corresponding protist cultures (Fig. 3A, Table S4), and nine of the enriched ASVs represented dominant culture genera identified in Fig. 1. A median 8.4% of the reads in each protist microbiome corresponded to ASVs enriched in the rhizosphere (0.8%-44%); over 17% of the microbiome associated with *Thaumatomonas*, *Flamella,* and *Allapsa* were rhizosphere-enriched. Some ASVs that were relatively rare in the protist microbiomes were also enriched, including *Brevundimonas* and an unclassified Burkholderiaceae. Many highly abundant protist-associated microbiome members, such as *Variovorax* (Fig. 1), were not enriched in the rhizosphere, indicating that high abundance in association with the protist was not predictive of rhizosphere enrichment.

**Figure 3:**
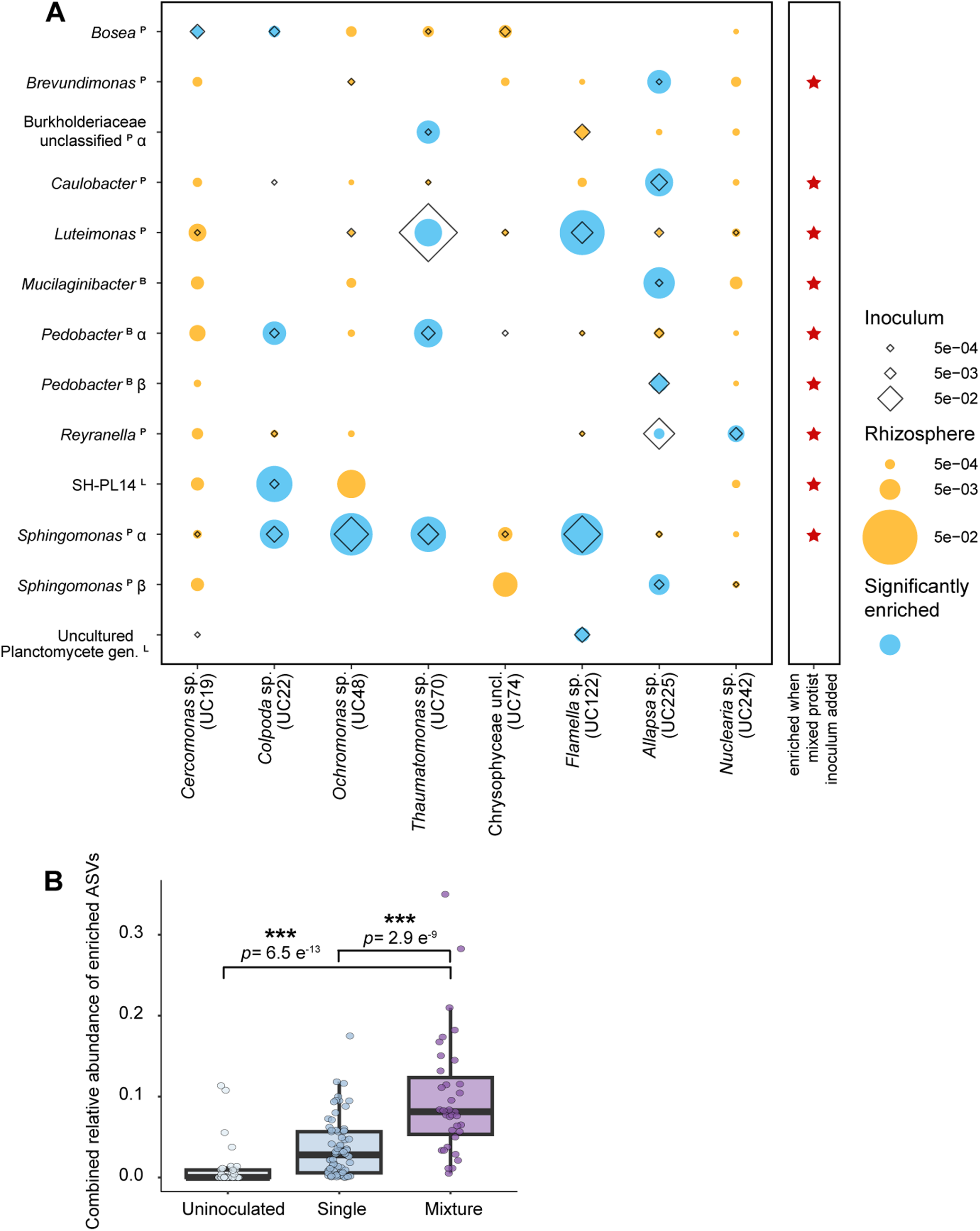
Enrichment of bacterial ASVs in the maize rhizosphere six weeks after protist inoculation onto germinated seeds. **A.** Enrichment by single protists. Diameter of diamonds and dots indicates the mean relative abundance of ASVs in the protist culture microbiome (n = 1) and the rhizosphere samples (n = 9 or 10), respectively. Blue color indicates statistical enrichment in the protist treatment relative to protist-uninoculated controls, as determined by DeSeq2. Red stars indicate that the bacteria were enriched when the eight protists were inoculated as a consortium. **B.** The proportion of rhizosphere reads representing protist-enriched ASVs after plants were inoculated with single protist cultures or mixed protists, or uninoculated. Asterisks indicate significant differences from the mixed protist treatment using Dunnett’s Multiple Comparison test (*** = p<0.0005).

Eight (61%) of the enriched bacterial ASVs were enriched by only one protist treatment. In contrast, a *Sphingomonas* ASV (*Sphingomonas-*α) was broadly enriched by four of the eight protist cultures and comprised up to 14.3% of the rhizosphere microbiome after inoculation. *Sphingomonas-*α was detected in all other protist microbiomes except that of *Nuclearia*. Collectively, these results demonstrate that a substantial proportion of the rhizosphere protist microbiome can colonize the rhizosphere, even after prolonged laboratory cultivation of protist isolates.

### Inoculating with multiple protists increased the abundance of protist-associated bacteria in the rhizosphere

Nine of the thirteen protist-enriched bacterial ASVs were also enriched in the rhizosphere after inoculation with an eight-protist mixture (Fig. 3A), indicating that inoculating diverse protist cultures has an additive effect in enriching protist-associated bacteria. The abundance of eight protist-associated bacterial taxa did not significantly differ whether single or mixed protists were inoculated (Fig. S2), even though the mixture contained a one-eighth dilution of each culture. However, the more diverse inoculum had a negative effect on the relative abundances of *Sphingomonas*-α, *Mucilaginibacter*, unclassified Burkholderiaceae sp., and two uncultured Planctomycetales sp., which were significantly lower in the mixed than single treatments (Fig. S2). Mixed protist inoculation resulted in an increased overall abundance of protist-associated bacteria in the rhizosphere (Fig. 3B). After inoculation with mixed protists, the thirteen protist-enriched bacterial ASVs comprised a median 9.9% of the bacterial reads in the rhizosphere, compared with 3.5% of reads after inoculation with single protists.

### Treatment with some protist cultures benefited root growth when inoculated individually, but not in mixture

We previously observed that plants inoculated with an antibiotic-treated protist culture had reduced root biomass compared with plants treated with a microbiome-replete protist (19), indicating that protist-associated bacteria were required to increase root growth. Therefore, we hypothesized that inoculation with mixed protist cultures would have a greater biomass effect than individual cultures due to the greater number and diversity of protist-associated bacteria enriched on the root. Among all data combined, plants inoculated with single protists had greater root:shoot biomass ratio than uninoculated plants, and showed a nonsignificant trend toward greater root biomass, but plants inoculated with the protist mixture did not (Fig. 4A-B). Among single protist treatments, *Thaumatomonas* culture-treated plants had increased root:shoot biomass ratios compared to the negative control, and *Flamella* and *Allapsa* treatments had increased root biomass (Fig. 4C-D). *Flamella* and *Allapsa* had the greatest bacterial richness among the protist cultures cultures (Table S1) and with *Thaumatomonas* had the greatest abundance of rhizosphere-enriched bacteria (Table S4), suggesting that bacterial richness or dominance of rhizosphere bacteria in the protist microbiome could be associated with greater biomass impact. However, no significant biomass phenotype was observed in plants treated with the protist mixture (Fig. 4C-D). Hence, the increased microbiome effect of the protist mixture did not translate to an increased plant biomass effect, and instead may have precluded the growth effects associated with *Thaumatomonas*, *Flamella*, and *Allapsa* treatments. Given the complex interactions among protists and bacteria, the taxonomic composition of enriched bacteria, rather than the overall richness or proportion of rhizosphere colonizers, may be more important for plant growth.

**Figure 4:**
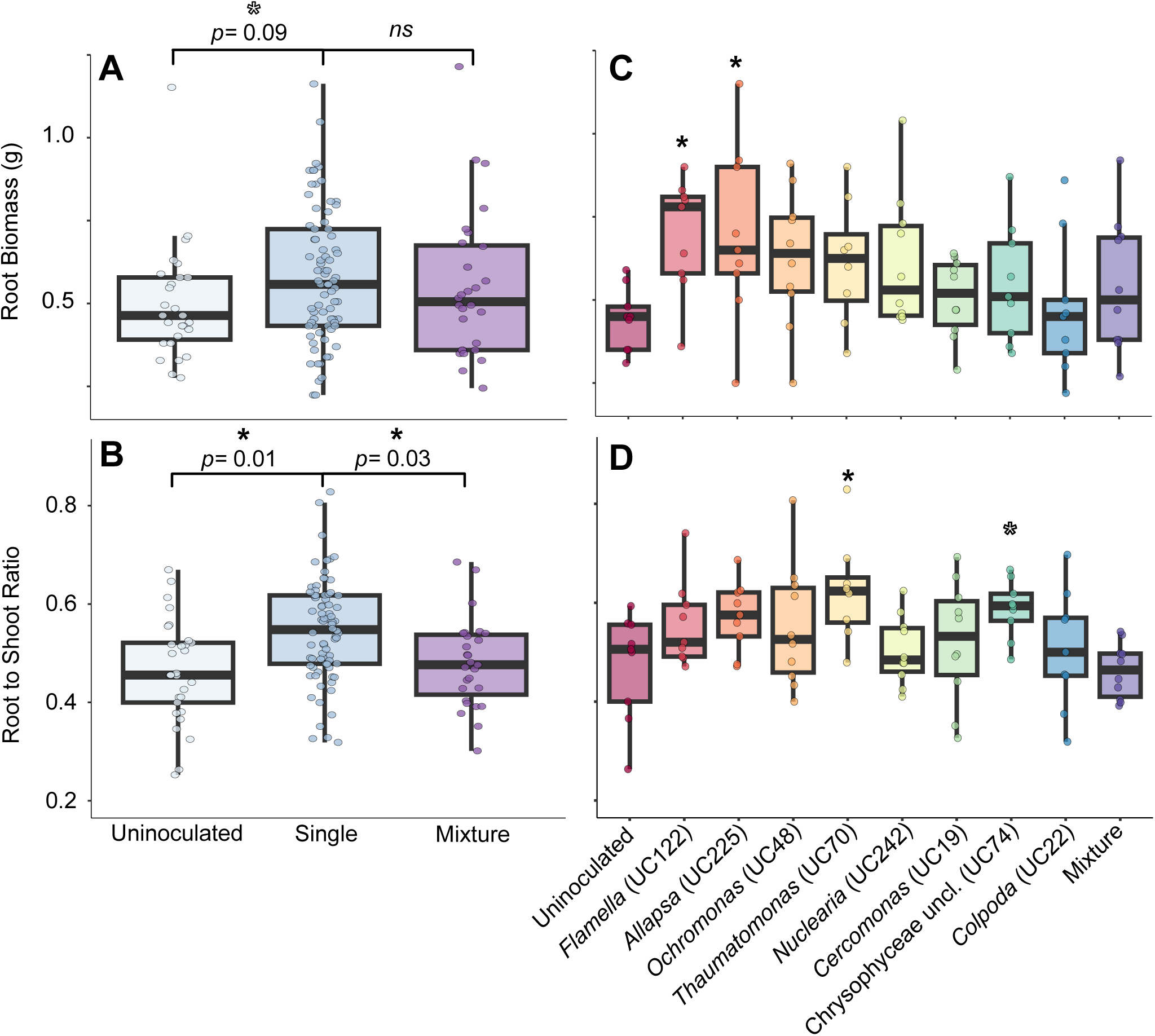
Biomass traits of maize plants inoculated with single protists and an eight-member consortium in a greenhouse study. **A-B.** Root biomass (**A**) and root-shoot mass ratio (**B**) six weeks after inoculation with zero (n = 29), one (n = 78), or a mixture of eight protists (n = 29) onto germinated maize seeds. Filled asterisks indicate significant differences from the single protist treatment at p < 0.05. **C-D.** Root biomass (**C**) and root-shoot mass ratio (**D**) after individual protist inoculation in the presence of protist-free soil bacteria (n = 10 for all treatments except UC22 and UC74, which had one plant fail to grow). Asterisks indicate significant differences from the no-protist treatment at p < 0.05.

### Protist inoculation was less impactful in the presence of a rhizosphere bacterial community

To determine whether the microbiome effect of protists was influenced by the surrounding bacterial community, we co-inoculated protist mixtures with bacterial communities cultured from three soil sources. Bacterial enrichment patterns following protist inoculation were similar among plants treated with bulk soil communities or no bacterial treatments; seven ASVs were enriched in multiple treatments (Fig. 5A). However, when protists were co-inoculated with rhizosphere bacterial community obtained from mature maize plants (RSC), only one ASV was enriched. We hypothesized that the rhizosphere soil community was a source of some protist-associated bacteria, and therefore they were not enriched by the protist inoculations. By comparing RSC-inoculated plants with controls, we confirmed that inoculation with RSC was sufficient to increase the rhizosphere abundance of four protist-associated ASVs in the rhizosphere (Fig. S3A). RSC-treated plants also had a higher combined abundance of protist-enriched taxa than bulk soil communities, and protist inoculation of RSC-treated plants did not increase the total abundance of protist-associated bacteria their abundance as observed in other treatments (Fig. S3B). This supports our previous finding that protist inoculation may substitute for rhizosphere soil in enriching certain bacterial taxa in the environment (19).

**Figure 5:**
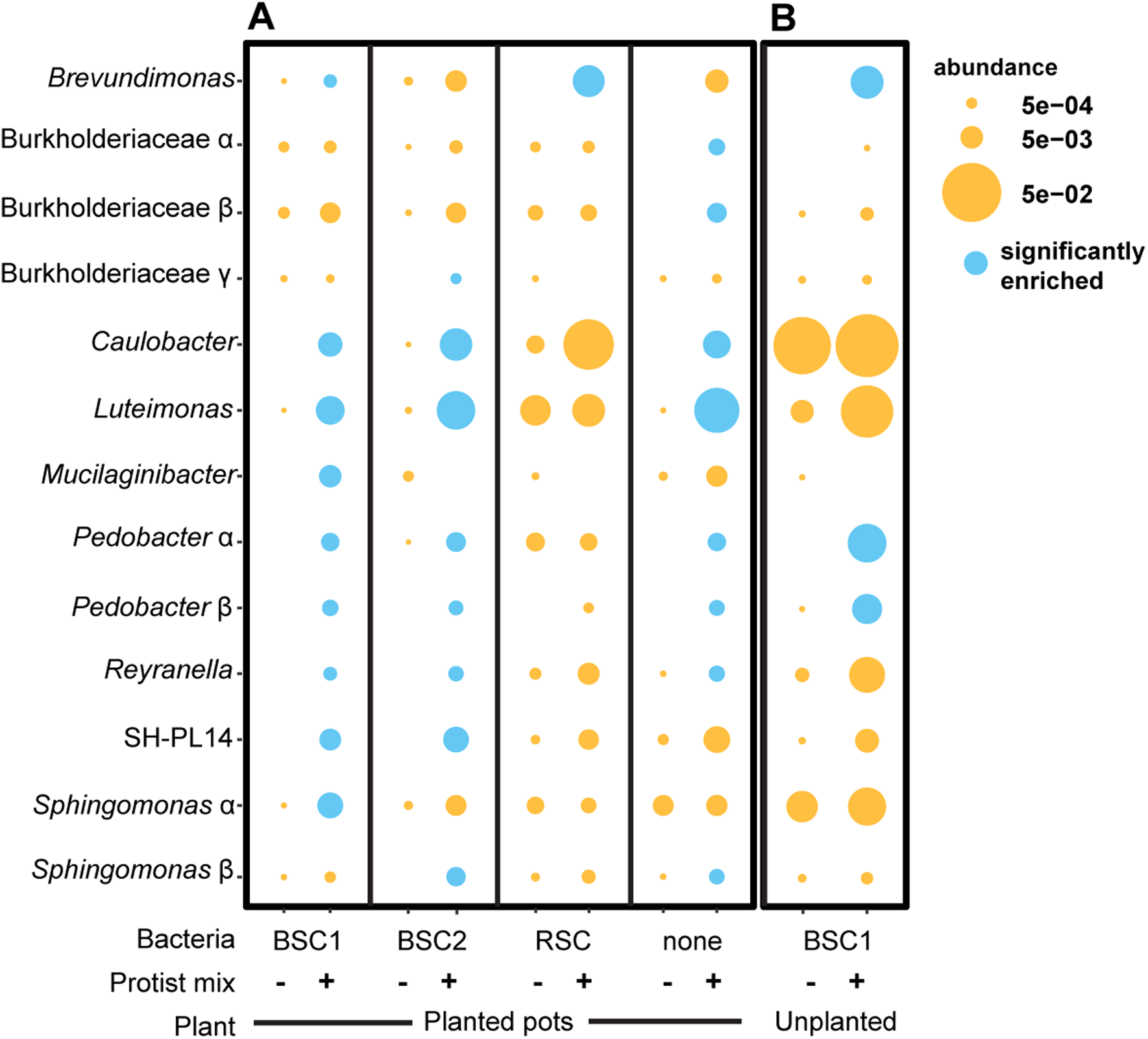
Effect of bacterial source (**A**) and rhizosphere environment (**B**) on protist enrichment of soil bacteria. **A.** The eight-protist consortium was co-inoculated with bulk soil bacteria from Lockwood Farm (BSC1), Griswold Farm (BSC2), maize rhizosphere soil bacteria from Lockwood Farm (RSC), or no bacteria. **B.** Enrichment after the eight-protist consortium was inoculated into the center of unplanted pots. Dot diameter indicates the mean relative abundance of the ASV (n = 9 or 10). Blue color indicates statistical enrichment in the protist mixture treatment relative to protist-uninoculated controls within each column.

We asked whether the rhizosphere environment was required for enrichment of protist-associated bacteria, or if protists alone were sufficient to enrich their abundance in soil. In unplanted pots, only *Brevundimonas* and *Pedobacter* ASVs showed increased abundance after protist inoculation, while others were only enriched in the presence of the plant (Fig. 5B). This result indicates that the protist-associated bacteria are specifically enriched by the rhizosphere. Unclassified Burkholderiaceae α was not consistently detected in unplanted pots (Fig. 5B, Fig. S3C), indicating that this taxon may obligately require either plant or protist hosts for environmental persistence.

### Seventeen additional bacterial ASVs were abundant in both rhizosphere soil and protist cultures

We detected diverse rhizosphere bacterial communities on both inoculated and untreated plants (Fig S1, Table S5), which could mask potential enrichment of some bacteria by protist inoculation. To determine whether other bacterial taxa colonized both rhizosphere and protist microbiomes, we identified ASVs in high rank abundance in both protist culture (top 20) and rhizosphere samples of the plants inoculated with the corresponding single protist (top 100). Thirty ASVs met these criteria, of which 17 had not been enriched by protist inoculation (Table 1, Table S6). Shared abundant ASVs included representatives of dominant genera that were abundant in protist cultures, including *Azospirillum*, *Pseudomonas*, and *Bradyrhizobium*. Notably, a *Variovorax* sp. ASV that was highly abundant in most protist microbiomes was also usually among the top 20 most abundant taxa on maize (Table S6).

**Table 1.** Protist culture microbiome bacterial ASVs that were enriched or abundant in the rhizosphere after protist inoculation.

| ASV<sup>a</sup> | Genus | Protist culture ID | Enriched in rhizosphere<sup>b</sup> | Abundant in rhizosphere<sup>c</sup> |
| --- | --- | --- | --- | --- |
| 1 | *Azospirillum* sp. | UC19, UC48, UC74, UC122 |  | X |
| 2 | *Bosea* sp. | UC74 |  | X |
| 3 | *Bosea* sp. | UC19, UC22, UC74 | X | X |
| 4 | *Bradyrhizobium* sp. | UC19, UC22, UC48, UC70, UC74, UC122 |  | X |
| 5 | *Brevundimonas* sp. | none | X |  |
| 6 | *Brevundimonas* sp. | UC242 |  | X |
| 7 | Burkholderiaceae uncl. | UC22 |  | X |
| 8 | Burkholderia uncl. | UC74 |  | X |
| 9 | Burkholderiaceae uncl. | UC122 | X | X |
| 10 | *Caulobacter* sp. | UC225, UC242 | X | X |
| 11 | *Caulobacter* sp. | UC19, UC225 |  | X |
| 12 | *Cupriavidus* sp. | UC225 |  | X |
| 13 | Enterobacteriaceae uncl. | UC48 |  | X |
| 14 | *Luteimonas* sp. | UC70, UC122 | X | X |
| 15 | *Mucilaginibacter* sp. | UC19, UC22 |  | X |
| 16 | *Mucilaginibacter* sp. | none | X |  |
| 17 | *Paenarthrobacter* sp. | UC242 |  | X |
| 18 | *Pedobacter* sp.-α | UC22, UC70 | X | X |
| 19 | *Pedobacter* sp.-β | UC225 | X | X |
| 20 | *Pseudomonas* sp. | UC19, UC48, UC74 |  | X |
| 21 | *Pseudoxanthomonas* sp. | UC122, UC19 |  | X |
| 22 | *Reyranella* sp. | UC225, UC242 | X | X |
| 23 | *Reyranella* sp. | none |  | X |
| 24 | *Rhodococcus* sp. | UC19, UC22, UC48, UC74, UC122, UC242 |  | X |
| 25 | *Sphingomonas* sp.-α | UC22, UC48, UC70, UC122 | X | X |
| 26 | *Sphingomonas* sp.-β | none | X |  |
| 27 | uncl. Planctomycetales | UC122 | X | X |
| 28 | uncl. Xanthobacteraceae | UC48, UC70, UC122, UC225 |  | X |
| 29 | uncultured gen. SH-PL14 | none | X |  |
| 30 | *Variovorax* sp. | UC19, UC22, UC48, UC70, UC74, UC122, UC242 |  | X |

Together, ASVs that were enriched or abundant on plants comprised a median 46.8% of the reads in the protist culture microbiomes (Table S4, range 6.6%-67.8%). This shows that several rhizosphere protist cultures were dominated by candidate rhizosphere-colonizing bacteria, including a wider range of taxa than was enriched by protist inoculation.

### Rhizosphere-enriched bacterial genera may promote or suppress protist growth

We lastly wondered why many rhizosphere colonizing bacteria are able to persist for long periods in culture with diverse types of protists. We hypothesized that the bacteria are not a food source for the protists, or alternatively, that they could provide a benefit to the protists. To test this, we isolated four bacteria from the protist cultures having 16S rRNA gene similarity to rhizosphere-enriched ASVs *Sphingomonas*-α (100% similarity), *Pedobacter*-β (98.4%), *Caulobacter* (98.0%), and *Mucilaginibacter* (96%). These genera were all enriched in the rhizosphere after inoculation with *Allapsa* (Fig. 3), which had a positive effect on root biomass. We measured the effects of these bacterial isolates on antibiotic-treated cultures of *Allapsa* as well as two other isolates from phylum Cercozoa, *Cercomonas* and *Thaumatomonas*, in which the four bacterial genera were detected (Fig. 1) but variably enriched (Fig. 3). All protists grew when cultures were supplemented with *Caulobacter* or *Sphingomonas,* but not when *Pedobacter, Mucilaginibacter,* or no bacteria were provided (Fig. S4); this indicates that *Pedobacter* and *Mucilaginibacter* isolates were not a source of prey (15). To determine whether the bacterial isolates suppress growth, we measured the effects of the bacteria on protists supplemented with an excess of inert prey (heat-killed *E. coli*). At 3 days, *Sphingomonas* increased growth of all three protists, increasing density of the cultures by 2.4 to 3.3-fold compared to *E. coli* alone. *Caulobacter sp.* had a strong positive effect on growth of *Allapsa* (2.4x, p<0.001), but a smaller effect on *Cercomonas* (1.5x, p=0.066), and no effect on *Thaumatomonas* (Fig. 6). Conversely, *Pedobacter* and *Mucilaginibacter* were strongly antagonistic to *Allapsa* growth, had differing positive and negative effects on *Cercomonas*, and had no significant effect on *Thaumatomonas*. Taken together, the results show that protist-associated rhizosphere bacteria use differing strategies to persist under predation pressure, supporting or inhibiting protist growth in broad or isolate-specific ways.

**Figure 6:**
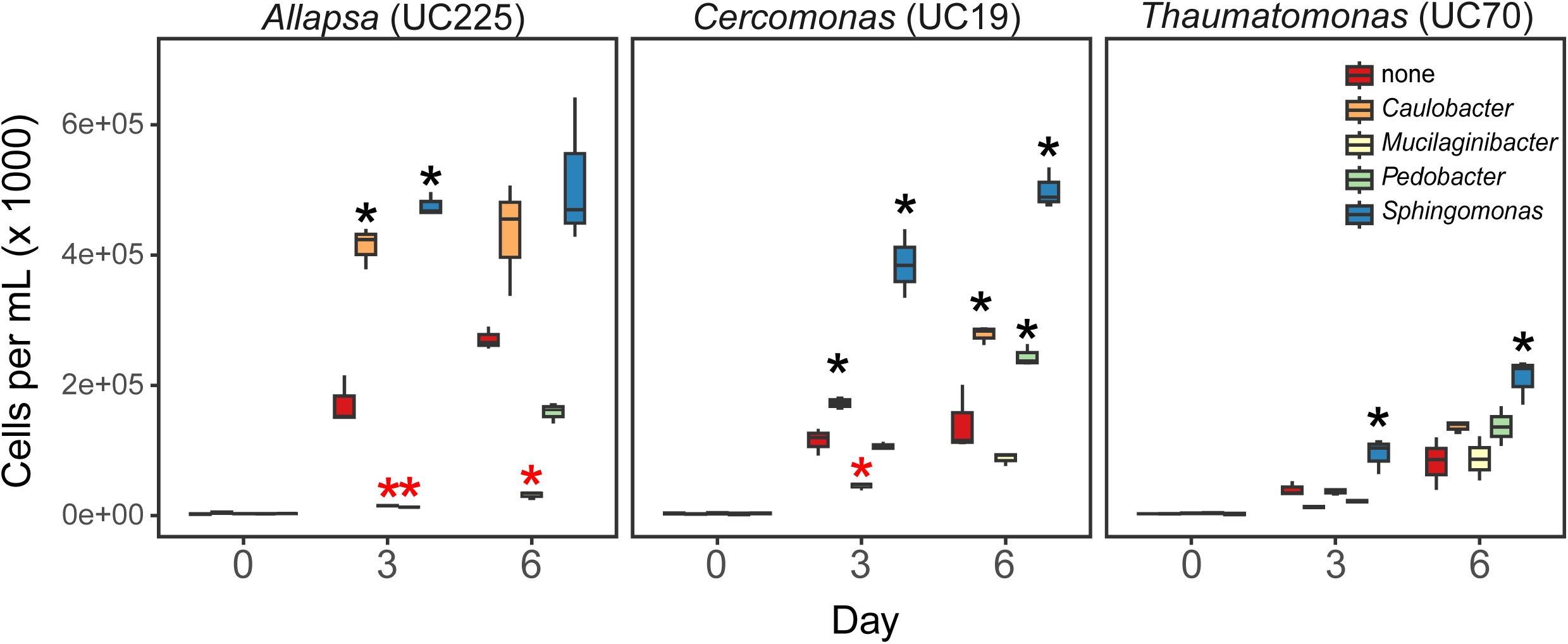
Cell counts of three protist taxa (*Allapsa*, *Cercomonas*, and *Thaumatomonas*) after being fed with heat-killed *E. coli* alone, or with heat-killed *E. coli* in addition to one of four bacterial cultures isolated from protist cultures (n = 3 for each treatment). Cells were counted at 0, 3 and 6 days after the cells were initially fed. Black asterisks indicate significantly more protists than those fed with heat-killed *E. coli* alone, while red asterisks indicate significantly fewer (p < 0.05).

## Discussion

Green algae in soil recruit bacterial communities that have strong structural and functional parallels to plant root communities (16), and the algal microbiome is a proposed source of plant beneficial bacteria (22). By surveying the microbiomes of diverse protist cultures and tracking their fates in the rhizosphere, this study demonstrates that a wide diversity of rhizosphere microeukaryotes outside the Chloroplastida have microbiomes largely comprised of plant-colonizing bacterial taxa. Diverse rhizosphere bacteria persisted for years in active cultures of protists from SAR, Excavata, and Amoebozoa supergroups, and maintained the capacity to colonize roots in a complex microbial environment. The findings support the hypothesis that, in addition to their roles imparting selective and behavioral advantages to rhizosphere bacteria (23, 24), predatory soil protists could recruit or promote a microbiome that contributes to rhizosphere establishment. Understanding how these bacteria interact with protists could help identify defense and signaling mechanisms important to bacterial survival, and may ultimately help enable the design of improved microbial consortia to benefit agriculture.

Profiling the protist microbiomes revealed broad taxonomic overlap in among protists independently isolated from different plants and field sites, indicating the broad prevalence of Burkholderiales and Hyphomicrobiales. They also included groups previously documented to survive protist internalization: members of *Sphingomonas*, *Burkholderia*, *Pseudomonas*, *Pseudoxanthomonas*, *Pedobacter*, *Mesorhizobium, and Acidovorax* have been identified in protist cells (Karaś et al., 2015; Lin et al., 2023; Morón et al., 2024; Zhang et al., 2023). *Variovorax* spp. are common epibionts and endosymbionts of protists (28, 29), and *Bosea* and *Reyranella* sp. are free-living protist mutualists that can be isolated using a strategy of protist co-culture (30, 31). While these studies support protist selection of these taxa, it’s also important to note that protist microbiome composition is likely shaped in part by the local soil microbiome, stochastic events during isolation, culture conditions, and bacteria-bacteria interactions. The microbiome communities identified here likely only represent a portion of true protist interactors in soil, and culture conditions could enhance associations that are rare or weak in nature. More work is needed to confirm which features of protist microbiomes represent replicable selection patterns and true symbioses. Enriched bacteria included two ASVs associated with uncultured Planctomyces species, indicating that protist inoculation could be a means for manipulation and rhizosphere introduction of some unculturable bacteria on the plant. However, we did not observe rhizosphere enrichment of the foodborne human pathogens hypothesized to be vectored by amoebae on produce (32, 33).

Protist-associated bacteria were enriched in the maize rhizosphere after six weeks, persisting even after the corresponding protists could no longer be detected. Some enriched genera identified here were previously correlated with protist abundance in plot studies, indicating that protist-bacterial network connections might reflect microbiome associations that are robust in the field. For example, *Luteimonas*, *Brevundimonas*, *Pseudomonas* and *Pseudoxanthomonas* were positively correlated with a *Colpoda* sp. in tomato rhizosphere (34). In this study the ASV *Sphingomonas-α* was enriched to high levels by the widest spectrum of protists; we previously identified the same ASV as the largest protist network hub on several field grown solanaceous crops (35). *Sphingomonas* abundance was previously linked to increased plant growth following *Cercomonas* inoculation on cucumber seedlings, and *Cercomonas* triggered biofilm formation of the bacteria *in vitro* (36). Our own observations that *Sphingomonas-α* is correlated with protists in the field, found in microbiomes of diverse protist cultures, enriched in the rhizosphere after protist inoculation, and broadly promotes protist growth *in vitro*, suggest that *Sphingomonas* could be a partner in widespread synergistic associations with protists and plants.

The effects of protist diversity on microbial communities is variable, and can shift dynamically under different environmental and biotic stress conditions (37, 38). We observed that inoculation with a mixture of protist cultures enriched a greater number and abundance of rhizosphere bacteria compared with single protist inoculation, but only a few single protists affected biomass traits. The increase in inoculum diversity may have affected the activity and distribution of bacteria through intermicrobial interactions, resulting in decreased abundance of potentially beneficial bacteria such as *Sphingomonas* and *Mucilaginibacter* (Fig. S1). Even when reducing inoculum diversity, changing community composition greatly shapes protist effects on the microbiome. We previously found that an 18-protist mixture had a biocontrol effect and greatly enriched the dominance of the genus *Duganella* in the rhizosphere after 3 weeks (19). The eight-protist mixture tested here did not enrich *Duganella*, which we later determined is found specifically in the microbiome of an *Ochrophyta* sp. isolate that was excluded from this study (UC29, Patel *et al*, in preparation). As we observed in this study, inclusion of a protist isolate hosting even a few rhizosphere-colonizing bacteria can have a significant impact on the microbiome. The observation of parallels among plant and protist microbiomes raises many questions: What is the mechanism and nature of protist interactions with its microbiome? Do protists provide the bacteria a fitness or other plant colonization advantage in the rhizosphere, or induce the bacteria to benefit the plant? The interactions identified here provide a starting point for understanding the ecology and agricultural significance of protist holobionts on plants.

## Methods

### Culture preparation

We selected protist cultures from a collection isolated from maize roots in 2020 (39). Protist cultures were maintained in PAGE’s amoebal saline (i.e., PAGE’s solution; 0.12 g of NaCl, 0.004 g of MgSO4·H2O, 0.004 g of CaCl2, 0.142 g of Na2HPO4, and 0.136 g of KH2PO4 in 1 liter of dH2O) amended with heat-killed *E. coli* as previously described (19). Protists had been maintained with monthly passaging for two years at the time of DNA extraction and inoculation onto plants. To prepare cultures for plant inoculation, protists were transferred to sterile PAGE’s solution and fed with heat-killed *E. coli* weekly for one month, grown just until encystment, centrifuged for 30 minutes at 2000 x g, resuspended in 4 mL PAGE’s solution, and pelleted again for 15 minutes at 2,000 x g before resuspension in 500 uL PAGE’s solution. Protist concentration was determined by counting under a microscope, then protists were diluted in PAGE’s solution to a concentration of 1000 cells/mL. Cultures were mixed in equal proportion to create the eight-protist mixture.

To generate protist-free soil bacteria, we collected 10 cm deep soil cores from five random locations in maize fields from Lockwood Farm (Hamden, CT) and Griswold, CT. We suspended 10 g of soil in 100 mL of sterile phosphate-buffered saline (PBS; pH 7.4; 8 g of NaCl, 1.44 g of Na2HPO4, 0.2 g of KCl, and 0.24 g of KH2PO4 in 1 liter of dH2O). Rhizosphere soil was collected from 5 plants of maize (*Zea mays* L.) inbred line B73 (stage V5) collected from the Lockwood field on the same day as the bulk soil. Roots were shaken to remove excess soil, and 5 roots per plant were agitated in sterile PBS to collect rhizosphere soil. Soil samples were plated in serial dilutions on both tryptic soya agar and Reasoner’s 2 agar plates (0.25 g of casein digest peptone, 0.25 g of peptic digest of animal tissue, 0.5 g of MgS04·7H2O, 0.3 g of C3H3NaO3, 0.5 g of casein acid hydrolysate, 0.3 g of K2HPO4, 0.5 g yeast extract, 0.5 g soluble starch, 0.5 g glucose, 15 g agar in 1 L of dH2O), which were incubated at 28°C for 72 hours. After incubation, the bacterial colonies from each plate were scraped and collected in PBS containing 20% glycerol and mixed by vortexing. Suspensions were stored at −80°C until use. One aliquot was used for colony enumeration. Bacterial inoculum for plant experiments was then prepared by thawing the tubes on ice and resuspending to 10^6^ cells/mL in sterile PBS.

### Plant inoculation and growth

We prepared a soil mixture containing 53% sand, 37% vermiculite, and 10% field soil from Lockwood farm, Hamden CT, USA, which is the same field from which all but two the protists were originally isolated. The mixture was wetted and autoclaved twice in 12 cm pots. Seeds of B73 maize were surface sterilized in 3% NaOCL for 1 min, then washed 10 times in sterilized dH2O. The seeds were germinated on sterile filter paper in the dark at 28°C for 2 days. In all but 20 pots, germinated seeds were placed into the middle of the pots at a depth of 4 cm and inoculated with 1 mL of protist inoculum (at a concentration of 10^3^ cells/mL) or sterile PAGE’s solution, then with 1 mL of bacteria (10^6^ cells/mL) or sterile phosphate buffered saline (pH 7.4), depending on the experimental treatment. Germinated seeds were inoculated by slowly pipetting the solution directly onto the emerging radicle before covering with the soil mixture. No germinated seeds were placed in the remaining 20 pots; instead, soil was removed to 4 cm in the center of the pot, and the soil in this location was inoculated with 1 mL of bulk soil bacteria and 1 mL of mixed protist inoculum (10 pots), or 1 mL of bulk soil bacteria alone (10 pots), before being covered by soil. Pots were arranged in a randomized block design and maintained in the greenhouse for six weeks during June and July, 2021 (16h days, temperatures 25-30°C). Plants were watered with autoclaved distilled water when pot weights decreased. At collection, roots were shaken vigorously to remove excess soil, and then placed in plastic bags containing 35 mL PBS. Roots were agitated for 30 seconds to remove attached soil, after which the rhizosphere suspensions were transferred to 50 mL tubes and stored at −80°C until DNA extraction. Root and shoot biomass were recorded after drying for one week at 50°C. For the plant-free replicates, soil was removed to a depth of 4 cm, and ∼1 g of soil was removed and transferred to a 50 mL tube containing 35 mL PBS.

### 16S and 18S rRNA gene library preparation

DNA extractions, PCR amplifications and sequencing was conducted following previously published protocols (19). The Illumina sequences can be found in NCBI GenBank under BioSample PRJNA950058.

### Read assembly and classification

We assembled the reads using mothur v. 1.44.0 (40). 16S contigs > 300 bp and 18S contigs < 90 bp or > 250 bp were removed, as were contigs containing ambiguous bases, or homopolymers greater than eight bases. Chimeras were removed in mothur using VSEARCH (41). 16S amplicon sequence variants (ASVs) were classified using the SILVA v. 132 database, while 18S ASVs were classified using the Protist Ribosomal Reference (PR^2^) v. 4.12 database (42). Both classifications used the naïve Bayesian classifier implemented in mothur. Libraries with fewer than 2000 reads were excluded from further analyses.

The sterilized PAGE’s solution sample was analyzed as a negative control. For the 16S analyses the control run totaled 7.9% reads of the next smallest library, while for the 18S analyses the control run totaled 7.7% reads of the next smallest library.

### Diversity and enrichment analyses

We calculated diversity metrics of inoculum samples and rhizosphere communities after subsampling to the smallest read number among inoculum samples or rhizosphere sample datasets using the phyloseq v. 1.38.0 package (43) implemented in R v. 4.1.2. The 16S communities were subsampled to 2195 reads for the protist inocula, 21547 for the bacterial inocula, and 2218 for the rhizosphere samples, while the 18S communities were subsampled to 14666 reads for the rhizosphere samples. Heatmap generation was performed using the MicEco v. 0.9.14 package (44), after removing ASVs with fewer than 5 reads in the dataset. Bacterial ASV enrichment between rhizosphere samples was performed using the DESeq2 v. 1.34.0 package (45) implemented in R. Adjusted P values for DESeq2 were calculated, and false positives were estimated using the Benjamini-Hochberg method.

### Bacterial isolation and identification

Protist culture solutions were diluted onto plates of TSA agar and incubated at 28° C for 3 days. 96 colonies were selected and cultured at random. Bacterial genomic DNA was extracted and amplified using the primers 27F (5’ AGAGTTTGATCCTGGCTCAG 3’) and 1492R (5’ GGTTACCTTGTTACGACTT 3’) (46).

Partial 16S sequences were sequenced using Sanger sequencing at the Keck DNA Sequencing Core at Yale University, New Haven, USA, and identified using Blast search in NCBI database. Strains identified as *Mucilaginibacter*, *Caulobacter*, *Sphingomonas*, and *Pedobacter* were respectively isolated from cultures of UC70, UC70, UC122, and UC225 and chosen for study. Sequence was deposited as Genbank accessions PQ741793-PQ741796.

### Protist growth assay

Cultures of *Allapsa* isolate UC225, *Cercomonas* isolate UC19, and *Thaumatomonas* isolate UC70 were treated with antibiotics to reduce endogenous bacteria as previously described (19). Protist cultures were grown in PAGE’s solution amended with heat-killed *E. coli* (O.D. 0.1) until the cells were freshly encysted. Cysts were centrifuged twice at 4000 g and resuspended in fresh PAGE’s solution to remove excess remaining killed *E. coli* cells, then enumerated and resuspended in PAGE’s solution to a concentration of 1000 cysts/mL. Bacterial cultures were grown for 72 h in Reasoner’s 2 broth media at 28°C.

Assays were conducted in 24-well cell culture plates. Protist cells were diluted to 100 cells in 1 mL of sterile PAGE’s solution, while bacteria were diluted to an O.D. of 0.07. In addition, wells were amended with heat-killed *E. coli* to an O.D. of 0.07. Three replicates each were set up of all protist-bacteria pairwise combinations, while three replicates each were set up of protists grown without added bacteria as controls. After the plates were set up, cells were visualized at 200 x magnification using a Zeiss ID02 Invertoscope inverted Microscope and photographed using an Axiocam 305 color camera.

Three photos were taken of each well in a triangular pattern. Cells were photographed every 24 hours for the following 6 days. Cells were enumerated using ImageJ, after which the number of cells in 1 mL were calculated for each well and timepoint.

### Statistical Analysis

Data were graphed using ggplot2 v. 3.4.0 (47), implemented in R version 4.3.0. Statistical differences between treatments and controls were determined using ANOVA followed by Dunnett’s multiple comparison test using the DescTools package in R. Cell count data were analyzed in R using a Dunnett Test to compare with the controls.

## Acknowledgements

This research was funded by AFRI Foundational Program grants from the United States Department of Agriculture National Institute of Food and Agriculture (2022-67013-37144 to L. R. Triplett, B. Steven, and S. J. Taerum). We thank Regan Huntley and Joseph Liquori for technical assistance.

## Supplementary figure captions

**Figure S1:** Effect of bacterial treatment on (**A**) the β-diversity of rhizosphere bacterial communities and (**B**) the biomass of maize plants. **A.** NMDS plot showing the clustering of 16S communities of maize rhizospheres after inoculation with bulk soil bacteria from Lockwood Farm (BSC1), Griswold Farm (BSC2), maize rhizosphere soil bacteria from Lockwood Farm (RSC), or no bacteria, and either no protists or the eight-protist consortium. Dashed lines represent 95% confidence intervals. **B.** Root biomass of maize plants six weeks after inoculation with the different bacterial treatments, with or without the eight-protist mixture.

**Figure S2:** Relative abundances of the 13 enriched bacteria in maize rhizospheres six weeks after inoculation with zero, one, or a mixture of eight protists onto germinated maize seeds. Asterisks indicate that relative abundances were significantly greater (p < 0.05) than in the mixture-inoculated plants based on a Dunnett’s Multiple Comparison Test.

**Figure S3:** Effect of rhizosphere on bacterial enrichment and abundance. **A.** Enrichment of five bacterial ASVs in maize rhizospheres inoculated with bulk soil bacteria from Lockwood Farm (BSC1) alone or maize rhizosphere soil bacteria from Lockwood Farm (RSC) alone compared with uninoculated controls. Circle diameters indicate mean relative abundances of bacteria. **B.** Proportion of reads of the 13 enriched bacterial ASVs in the plants inoculated with no protists, but different bacterial inocula. Asterisks indicate that the read proportions are significantly greater when plants were inoculated with protists compared with the plants that were not inoculated with protists at p < 0.005 (**) and p < 0.001 (***). **C.** Relative abundance of the bacterial ASV Burkholderiaceae unclassified β in soil inoculated with the protist mixture or no protist, with or without the plant.

**Figure S4:** Cell counts of three protist taxa (*Allapsa*, *Cercomonas*, and *Thaumatomonas*) after being fed with no bacteria, or one of four bacterial isolates from protist cultures (n = 3 for each treatment). Cells were counted at 0, 3 and 6 days after the cells were initially fed. Black asterisks indicate significantly more protists than those fed with no bacteria (p < 0.05) based on a Dunnett’s Multiple Comparison Test.

## References

1. Oliverio AM, Geisen S, Delgado-Baquerizo M, Maestre FT, Turner BL, Fierer N. 2020. The global-scale distributions of soil protists and their contributions to belowground systems. Sci Adv 6:eaax8787.

2. Gao Z, Karlsson I, Geisen S, Kowalchuk G, Jousset A. 2019. Protists: puppet masters of the rhizosphere microbiome. Trends Plant Sci 24:165–176.

3. Martins SJ, Taerum SJ, Triplett L, Emerson JB, Zasada I, de Toledo BF, Kovac J, Martin K, Bull CT. 2022. Predators of soil bacteria in plant and human health. Phytobiomes J 6:184–200.

4. Nguyen TBA, Chen Q-L, Yan Z-Z, Li C, He J-Z, Hu H-W. 2023. Trophic interrelationships of bacteria are important for shaping soil protist communities. Environ Microbiol Rep 15:298–307.

5. Xiong W, Song Y, Yang K, Gu Y, Wei Z, Kowalchuk GA, Xu Y, Jousset A, Shen Q, Geisen S. 2020. Rhizosphere protists are key determinants of plant health. Microbiome 8:27.

6. Yue Y, Liu C, Xu B, Wang Y, Lv Q, Zhou Z, Li R, Kowalchuk GA, Jousset A, Shen Q, Xiong W. 2023. Rhizosphere shapes the associations between protistan predators and bacteria within microbiomes through the deterministic selection on bacterial communities. Environ Microbiol 25:3623–3629.

7. Röttjers L, Faust K. 2018. From hairballs to hypotheses–biological insights from microbial networks. FEMS Microbiol Rev 42:761–780.

8. Cheng X-Y, Tian X-L, Wang Y-S, Lin R-M, Mao Z-C, Chen N, Xie B-Y. 2013. Metagenomic analysis of the pinewood nematode microbiome reveals a symbiotic relationship critical for xenobiotics degradation. Sci Rep 3:1869.

9. Guo H, Glaeser SP, Alabid I, Imani J, Haghighi H, Kämpfer P, Kogel K-H. 2017. The abundance of endofungal bacterium *Rhizobium radiobacter* (syn. Agrobacterium tumefaciens) increases in its fungal host Piriformospora indica during the tripartite sebacinalean symbiosis with higher plants. Front Microbiol 8.

10. Partida-Martinez LP, Hertweck C. 2005. Pathogenic fungus harbours endosymbiotic bacteria for toxin production. Nature 437:884–888.

11. Xu S, Liu Y-X, Cernava T, Wang H, Zhou Y, Xia T, Cao S, Berg G, Shen X-X, Wen Z, Li C, Qu B, Ruan H, Chai Y, Zhou X, Ma Z, Shi Y, Yu Y, Bai Y, Chen Y. 2022. *Fusarium* fruiting body microbiome member *Pantoea agglomerans* inhibits fungal pathogenesis by targeting lipid rafts. Nat Microbiol 7:831– 843.

12. Kjeldgaard B, Listian SA, Ramaswamhi V, Richter A, Kiesewalter HT, Kovács ÁT. 2019. Fungal hyphae colonization by *Bacillus subtilis* relies on biofilm matrix components. Biofilm 1:100007.

13. Yergaliyev T, Alexander-Shani R, Dimerets, H. 2020. Bacterial community structure dynamics in *Meloidogyne incognita*-infected roots and its role in worm-microbiome interactions. https://journals.asm.org/doi/epdf/10.1128/msphere.00306-20. Retrieved 4 September 2024.

14. Amin SA, Hmelo LR, Van Tol HM, Durham BP, Carlson LT, Heal KR, Morales RL, Berthiaume CT, Parker MS, Djunaedi B, Ingalls AE, Parsek MR, Moran MA, Armbrust EV. 2015. Interaction and signalling between a cosmopolitan phytoplankton and associated bacteria. Nature 522:98–101.

15. Steele MI, Peiser JM, Shreenidhi PM, Strassmann JE, Queller DC. 2023. Predation-resistant *Pseudomonas* bacteria engage in symbiont-like behavior with the social amoeba *Dictyostelium discoideum*. ISME J 17:2352–2361.

16. Durán P, Flores-Uribe J, Wippel K, Zhang P, Guan R, Melkonian B, Melkonian M, Garrido-Oter R. 2022. Shared features and reciprocal complementation of the *Chlamydomonas* and *Arabidopsis* microbiota. Nat Commun 13:406.

17. Hawxhurst CJ, Micciulla JL, Bridges CM, Shor M, Gage DJ, Shor LM. 2022. Soil protists can actively redistribute beneficial bacteria along *Medicago truncatula* roots. bioRxiv 10.1101/2021.06.16.448774.

18. Micciulla JL, Shor LM, Gage DJ. 2024. Enhanced transport of bacteria along root systems by protists can impact plant health. Appl Environ Microbiol 90:e02011–23.

19. Taerum SJ, Patel RR, Micciulla J, Steven B, Gage D, Triplett LR. 2024. Establishment and effect of a protist consortium on the maize rhizosphere. Phytobiomes J PBIOMES-04–24-0035-R.

20. Taerum SJ, Micciulla J, Corso G, Steven B, Gage DJ, Triplett LR. 2022. 18S rRNA gene amplicon sequencing combined with culture-based surveys of maize rhizosphere protists reveal dominant, plant-enriched and culturable community members. Environ Microbiol Rep 14:110–118.

21. Ceja-Navarro JA, Wang Y, Ning D, Arellano A, Ramanculova L, Yuan MM, Byer A, Craven KD, Saha MC, Brodie EL, Pett-Ridge J, Firestone MK. 2021. Protist diversity and community complexity in the rhizosphere of switchgrass are dynamic as plants develop. Microbiome 9:96.

22. Zheng Q, Hu Y, Kosina SM, Van Goethem MW, Tringe SG, Bowen BP, Northen TR. 2023. Conservation of beneficial microbes between the rhizosphere and the cyanosphere. New Phytol 240:1246–1258.

23. Dumack K, Feng K, Flues S, Sapp M, Schreiter S, Grosch R, Rose LE, Deng Y, Smalla K, Bonkowski M. 2022. What drives the assembly of plant-associated protist microbiomes? Investigating the effects of crop species, soil type and bacterial microbiomes. Protist 173:125913.

24. Yue Y, Liu C, Xu B, Wang Y, Lv Q, Zhou Z, Li R, Kowalchuk GA, Jousset A, Shen Q, Xiong W. 2023. Rhizosphere shapes the associations between protistan predators and bacteria within microbiomes through the deterministic selection on bacterial communities. Environ Microbiol 25:3623–3629.

25. Karaś MA, Turska-Szewczuk A, Trapska D, Urbanik-Sypniewska T. 2015. Growth and survival of *Mesorhizobium loti* Inside *Acanthamoeba* enhanced its ability to develop more nodules on *Lotus corniculatus*. Microb Ecol 70:566–575.

26. Lin C, Li L-J, Ren K, Zhou S-Y-D, Isabwe A, Yang L-Y, Neilson R, Yang X-R, Cytryn E, Zhu Y-G. 2023. Phagotrophic protists preserve antibiotic-resistant opportunistic human pathogens in the vegetable phyllosphere. ISME Commun 3:94.

27. Morón Á, Tarhouchi AE, Belinchón I, Valenzuela JM, de Francisco P, Martín-González A, Amaro F. 2024. Protozoan predation enhances stress resistance and antibiotic tolerance in *Burkholderia cenocepacia* by triggering the SOS response. ISME J wrae014.

28. Zhang X, Bi L, Gentekaki E, Zhao J, Shen P, Zhang Q. 2023. Culture-independent single-cell PacBio sequencing reveals epibiotic *Variovorax* and nucleus associated *Mycoplasma* in the microbiome of the marine benthic protist *Geleia* sp. YT (Ciliophora, Karyorelictea). Microorganisms 11.

29. Corsaro D, Michel R, Walochnik J, Müller K-D, Greub G. 2010. *Saccamoeba lacustris*, sp. nov. (Amoebozoa: Lobosea: Hartmannellidae), a new lobose amoeba, parasitized by the novel chlamydia ‘*Candidatus Metachlamydia lacustris*’ (Chlamydiae: *Parachlamydiaceae*). Eur J Protistol 46:86–95.

30. Pagnier I, Croce O, Robert C, Raoult D, La Scola B. 2012. Genome sequence of *Reyranella massiliensis*, a bacterium associated with amoebae. J Bacteriol 194:5698–5698.

31. Thomas V, Casson N, Greub G. 2007. New *Afipia* and *Bosea* strains isolated from various water sources by amoebal co-culture. Syst Appl Microbiol 30:572–579.

32. Vaerewijck M, Houf K. 2015. 4 - The role of free-living protozoa in protecting foodborne pathogens, p. 81–101. In Sofos, J (ed.), Advances in Microbial Food Safety. Woodhead Publishing, Oxford.

33. Moreno-Mesonero L, Hortelano I, Moreno Y, Ferrús MA. 2020. Evidence of viable *Helicobacter pylori* and other bacteria of public health interest associated with free-living amoebae in lettuce samples by next generation sequencing and other molecular techniques. Int J Food Microbiol 318:108477.

34. Guo S, Jiao Z, Yan Z, Yan X, Deng X, Xiong W, Tao C, Liu H, Li R, Shen Q, Kowalchuk GA, Geisen S. 2024. Predatory protists reduce bacteria wilt disease incidence in tomato plants. Nat Commun 15:829.

35. Taerum SJ, Steven B, Gage D, Triplett LR. 2022. Dominance of Ciliophora and Chlorophyta among ohyllosphere protists of solanaceous plants. Phytobiomes J PBIOMES-04-22-0021-FI.

36. Guo S, Geisen S, Mo Y, Yan X, Huang R, Liu H, Gao Z, Tao C, Deng X, Xiong W, Shen Q, Kowalchuk GA, Li R. 2024. Predatory protists impact plant performance by promoting plant growth-promoting rhizobacterial consortia. ISME J 18:wrae180.

37. Berlinches De Gea A, Li G, Chen JO, Wu W, Kohra A, Aslan SK, Geisen S. 2023. Increasing soil protist diversity alters tomato plant biomass in a stress-dependent manner. Soil Biol Biochem 186:109179.

38. Xiong W, Li R, Guo S, Karlsson I, Jiao Z, Xun W, Kowalchuk GA, Shen Q, Geisen S. 2019. Microbial amendments alter protist communities within the soil microbiome. Soil Biol Biochem 135:379–382.

39. Taerum SJ, Micciulla J, Corso G, Steven B, Gage DJ, Triplett LR. 2022. 18S _R_RNA gene amplicon sequencing combined with culture-based surveys of maize rhizosphere protists reveal dominant, plant-enriched and culturable community members. Environ Microbiol Rep 14:110–118.

40. Schloss PD, Westcott SL, Ryabin T, Hall JR, Hartmann M, Hollister EB, Lesniewski RA, Oakley BB, Parks DH, Robinson CJ, Sahl JW, Stres B, Thallinger GG, Van Horn DJ, Weber CF. 2009. Introducing mothur: open-source, platform-independent, community-supported software for describing and comparing microbial communities. Appl Environ Microbiol 75:7537–7541.

41. Rognes T, Flouri T, Nichols B, Quince C, Mahé F. 2016. VSEARCH: a versatile open source tool for metagenomics. PeerJ 4:e2584.

42. Guillou L, Bachar D, Audic S, Bass D, Berney C, Bittner L, Boutte C, Burgaud G, de Vargas C, Decelle J, del Campo J, Dolan JR, Dunthorn M, Edvardsen B, Holzmann M, Kooistra WHCF, Lara E, Le Bescot N, Logares R, Mahé F, Massana R, Montresor M, Morard R, Not F, Pawlowski J, Probert I, Sauvadet A-L, Siano R, Stoeck T, Vaulot D, Zimmermann P, Christen R. 2013. The Protist Ribosomal Reference database (PR2): a catalog of unicellular eukaryote Small Sub-Unit rRNA sequences with curated taxonomy. Nucleic Acids Res 41:D597–D604.

43. McMurdie PJ, Holmes S. 2013. phyloseq: An R package for reproducible interactive analysis and graphics of microbiome census data. PLoS ONE 8:e61217.

44. Russel J. 2021. Russel88/MicEco: v0.9.15. Zenodo.

45. Love MI, Huber W, Anders S. 2014. Moderated estimation of fold change and dispersion for RNA-seq data with DESeq2. Genome Biol 15:550.

46. Lane DJ. 1991. 16S/23S sequencing, p. 115–175. In Nucleic Acid Technologies in Bacterial Systematics. Wiley, New York, NY, U.S.A.

47. Wilkinson L. 2011. ggplot2: elegant graphics for data analysis by WICKHAM, H. Biometrics 67:678– 679.

