## Supplementary figures and images for "Rhizosphere-colonizing bacteria persist in the protist microbiome"

### Fig. S1

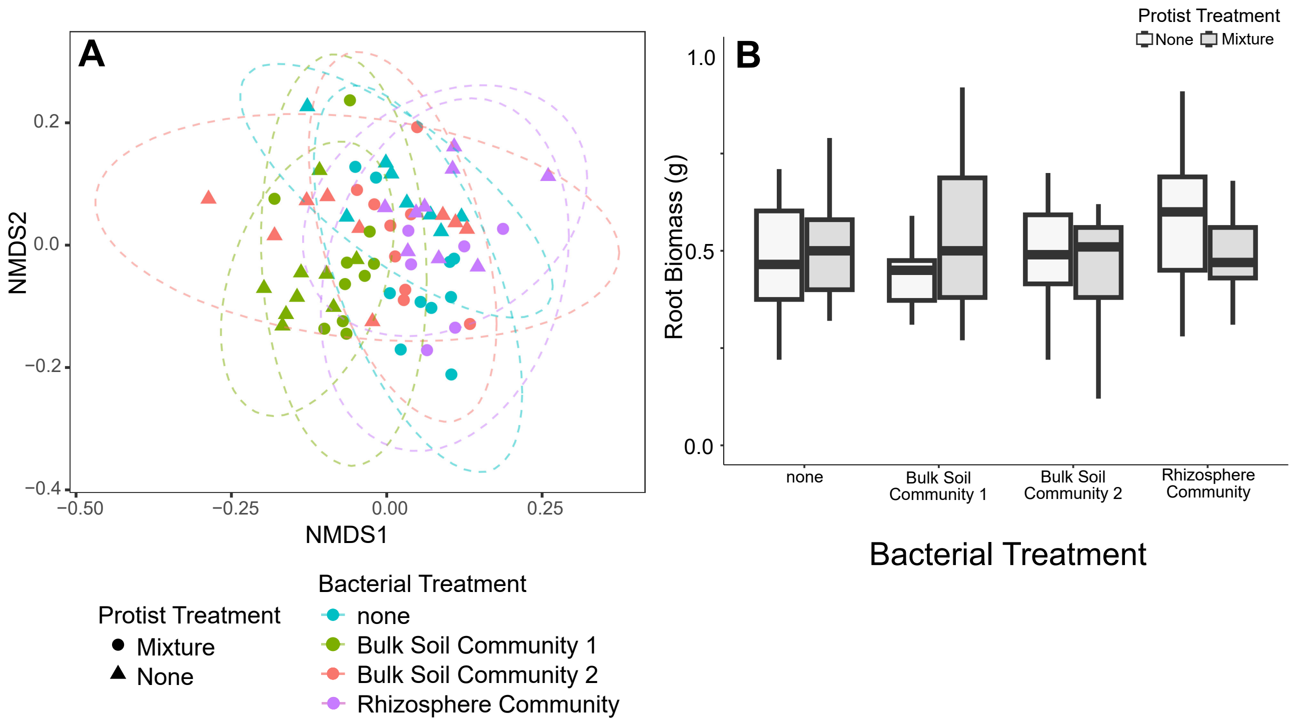

### Fig. S2

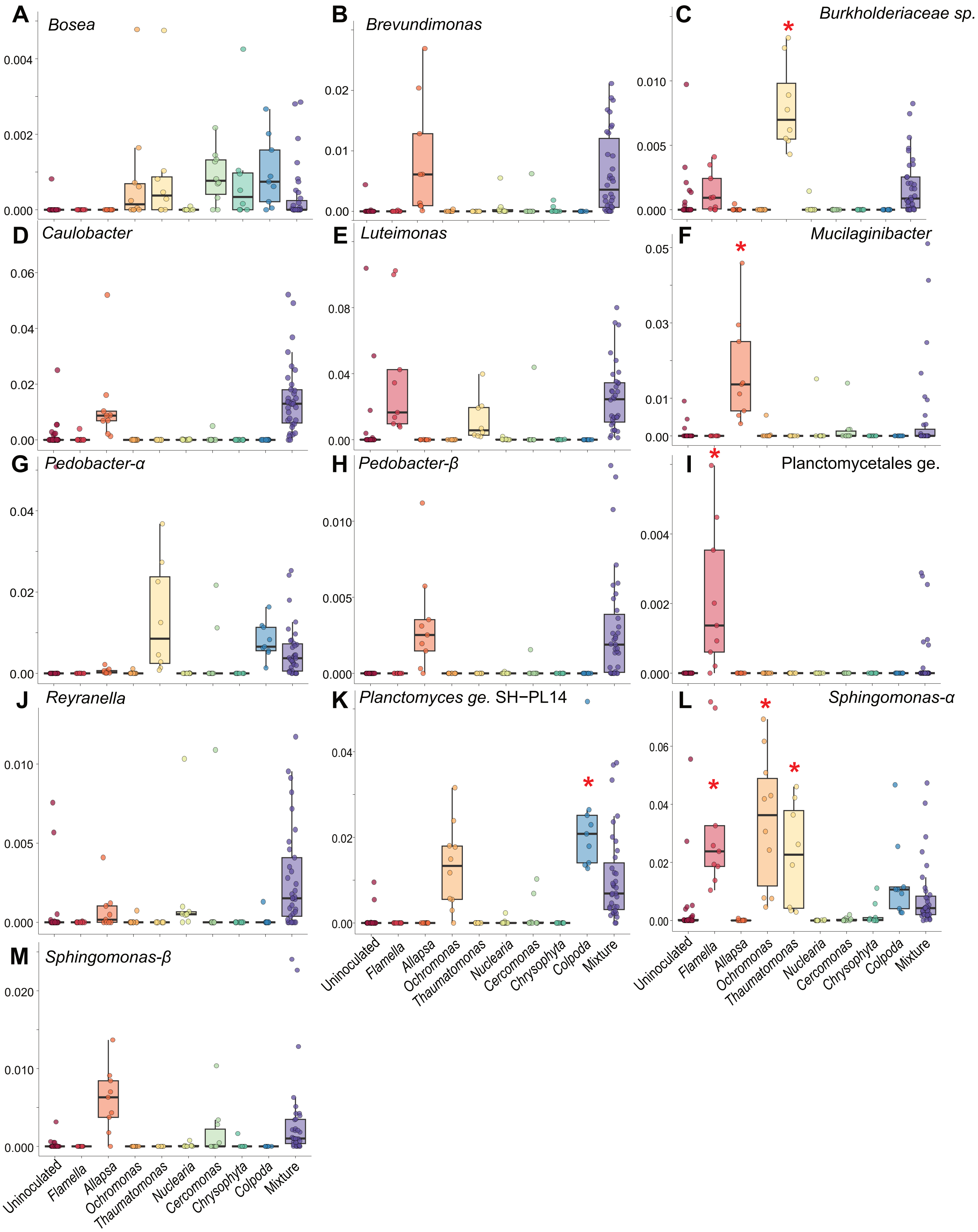

### Fig. S3

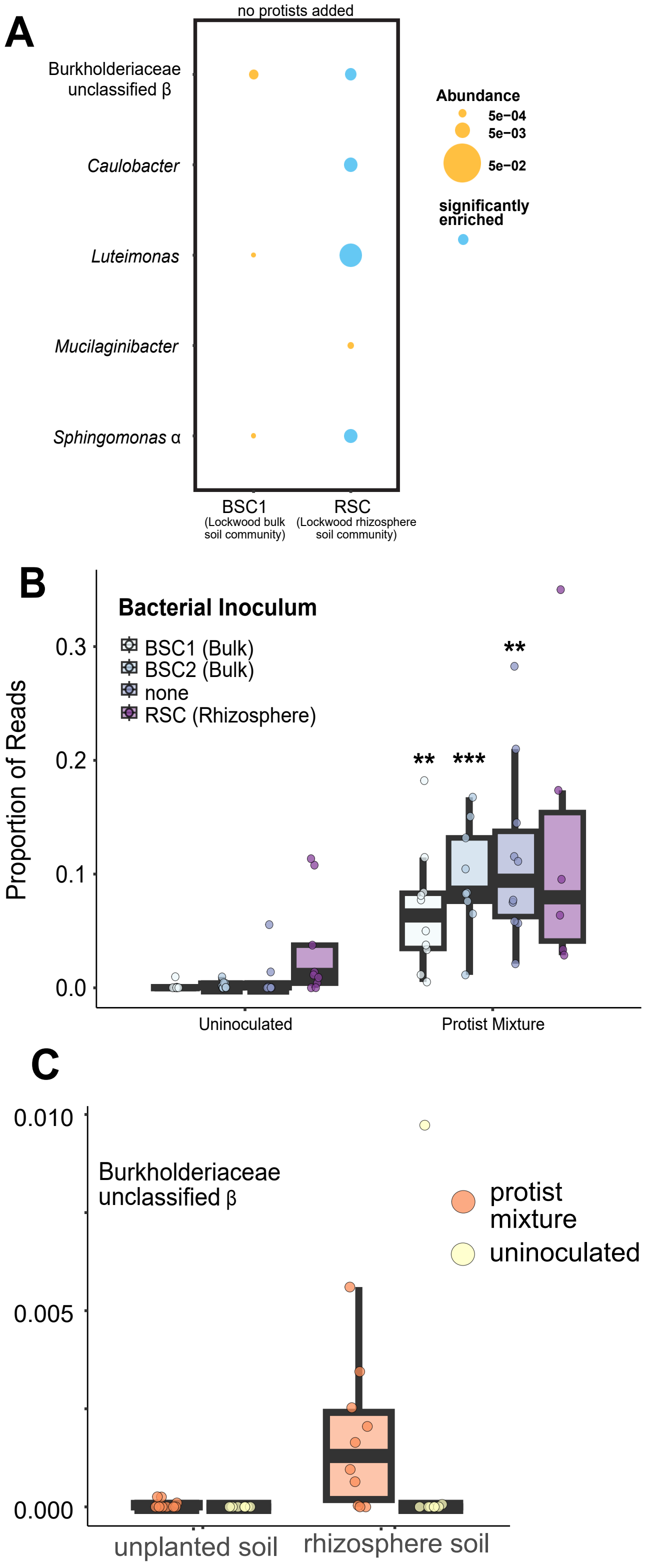

### Fig. S4

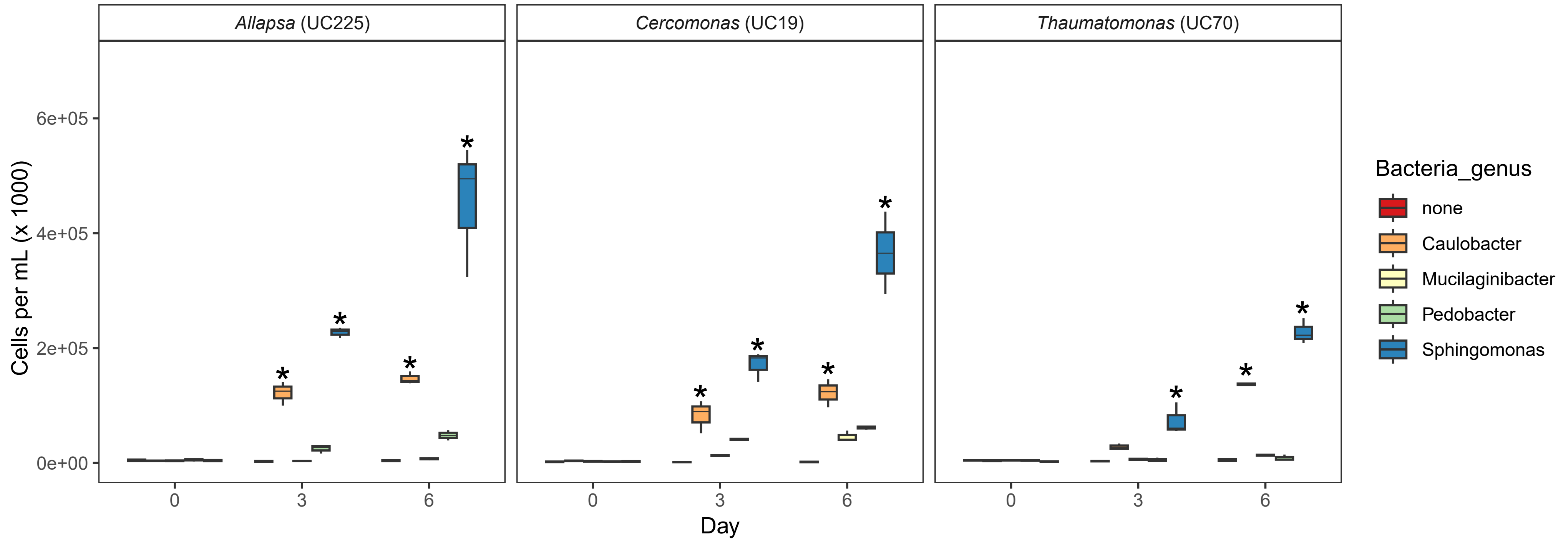
